# An inspectable Grover algorithm-based amplitude-amplification workflow for k-mer discovery and motif scanning with genomic coordinate traceability

**DOI:** 10.64898/2026.10.09.758025

**Authors:** Akshay Uttarkar, Vidya Niranjan

**Affiliations:** Department of Biotechnology, R V College of Engineering, (Affiliated with Visvesvaraya Technological University, Belagavi-590018), Bangalore, 560059, India; MIT Vishwaprayag University, Solapur, Maharashtra, India

**Keywords:** Quantum computing, amplitude amplification, Grover search, motif discovery, k-mers, genomics

## Abstract

Motif discovery and motif scanning remain core primitives in genomics, but both tasks face combinatorial growth as k-mer length increases and as mismatch-tolerant searches expand the candidate space. In this work, we present MotifQu, a Python package and command-line tool that maps two common motif-analysis workflows—(i) scanning a genome for a user-specified consensus motif and (ii) de novo discovery of recurrent k-mers above a user-defined count threshold—onto Grover-style amplitude amplification circuits executed reproducibly via Qiskit Aer statevector simulation.

We evaluate MotifQu on the SARS-CoV-2 reference contig (29,903 bp) and S. cerevisiae S288C reference genome, where the classical census identifies strongly non-uniform motif frequencies dominated by A/T-rich 5-mers and returns explicit coordinate lists across the full genome span.

We further execute the same logical 10-qubit search on IBM hardware showing that while end-to-end execution is feasible, transpilation-induced circuit depth and entangling-gate volume produce a diffuse measurement distribution whose probabilities remain close to the uniform 1/1024 scale, limiting observable amplitude concentration under current NISQ constraints.

Finally, we demonstrate the full discovery workflow on the S. cerevisiae S288C reference genome (12,157,105 bp) to illustrate search results (ranked motifs, probability summaries, circuit diagrams, and position plots) and to highlight the regime where a large, marked set reduces the sharpness of Grover amplification.

## Introduction

Motifs underpin many functional genomic elements, including transcription factor binding sites, promoters, and repeat structures. Practical workflows typically split into two tasks: (i) scanning a genome for occurrences of a known consensus motif, and (ii) discovering enriched short patterns (k-mers) that recur above background expectations [1]. MotifQu is a Python package and command-line tool that implements both tasks using Grover-style amplitude amplification [2] with circuits executed via Qiskit Aer’s statevector simulator [3, 4].

MotifQu provides two core subcommands: motifqu search for locating a user-specified motif with an optional Hamming-mismatch tolerance, and motifqu discover for identifying significant k-mers whose genome-wide counts exceed a user-defined threshold. In addition, MotifQu includes utilities for common regulatory-pattern workflows, such as expanding IUPAC-degenerate motifs into concrete sequences [5] and listing a curated set of known biological motif consensuses.

Motif discovery and motif scanning are foundational operations in computational genomics because many biological mechanisms are mediated by short, recurrent sequence patterns. Examples include transcription factor binding sites, promoter elements such as TATA-box-like patterns, and other regulatory signals that influence gene expression. In their simplest form, motifs can be represented as fixed strings (k-mers) or as degenerate consensuses (e.g., IUPAC codes), while more expressive models capture position-specific preferences (e.g., position weight matrices). Independent of the representation, the computational problem is driven by the same combinatorial structure: for DNA, there are 4*^k^* possible k-mers, and for a genome of length *G* there are *W* = *G* − *k* + 1 candidate windows to examine.

Classical bioinformatics provides mature and statistically principled solutions. For *de novo* motif discovery, the MEME Suite is widely used and implements expectation-maximization-based approaches that learn motifs from sets of sequences and provide downstream analysis utilities [6]. For scanning a known motif model across sequences, tools such as FIMO perform efficient searches and report statistically interpretable significance measures for predicted motif occurrences [7]. These tools remain the standard for production genomics because they scale to large datasets, support rich motif models, and integrate with established statistical workflows.

At the same time, motif-related tasks expose computational bottlenecks that become more pronounced as search spaces expand. Exhaustive enumeration of all k-mers scales as *O*(4*^k^*) in the number of candidate patterns, and genome-wide scanning scales as *O*(*W*) per motif under straightforward formulations. While classical accelerations (indexing, heuristics, vectorization, and statistical pruning) substantially improve practical performance, the underlying combinatorial growth remains a key driver of runtime and memory use, especially when exploring many candidate motifs, large mismatch neighborhoods, or multiple genomes.

Quantum algorithms are of interest here because they provide a different asymptotic scaling for unstructured search. Grover’s algorithm and related amplitude amplification techniques [8] can find marked items in an unstructured space of size *N* using *O(√N)* oracle queries, compared to *O*(*N*) for classical exhaustive search in the same oracle model [2,9]. In the motif context, the search space can correspond to (i) genomic windows when locating a given motif (with a marking oracle based on a mismatch predicate), or (ii) the k-mer space when identifying k-mers that exceed a chosen significance threshold (with a marking oracle based on precomputed counts). Although near-term quantum hardware and noise constraints mean that end-to-end quantum speedups for practical genomics are not yet established, transparent implementations are essential for studying where (and where not) amplitude amplification could help— particularly for separating the cost of classical preprocessing from the quantum query complexity of the amplified search.

Widely used motif-analysis tools such as the MEME Suite [6] and FIMO [7] implement statistically grounded discovery and scanning using classical algorithms, and remain the default choice for production genomics. However, the combinatorial nature of motif search spaces— particularly the 4*^k^* space of DNA k-mers—also makes motif analysis a natural target for quantum search primitives, providing a clear test-bed for quantum search primitives in genomics [2,9].

Classical motif analysis tools are mature and feature-rich: the MEME Suite [6] provides expectation-maximization-based *de novo* motif discovery, and FIMO [7] offers fast scanning with statistical significance estimates. In contrast, quantum approaches to bioinformatics remain nascent, and reproducible, end-to-end implementations that clearly separate (a) classical preprocessing from (b) quantum amplitude amplification are scarce.

This work introduces MotifQu, a small, reproducible research artifact that makes the mapping from motif analysis to Grover-style search explicit. Rather than attempting to replace established classical tools, MotifQu targets the methodological gap between theoretical proposals and practical experimentation: it provides inspectable oracle construction, clear reporting of genomic coordinates, and diagnostic outputs (circuit structure and probability summaries) needed to validate and benchmark quantum motif workflows under simulation.

This work aims to (i) implement motif search with an optional Hamming-distance tolerance by applying Grover-style amplitude amplification over candidate genomic windows, (ii) implement k-mer (motif) discovery by amplifying k-mers whose genome-wide counts meet or exceed a user-defined threshold, and (iii) prioritize transparency and reproducibility by exposing the marked set and oracle construction and generating interpretable outputs (hits with coordinates, probability-ranked motifs, and diagnostic plots) for benchmarking and teaching.

## Installation and usage

MotifQu is available on PyPI (Python Package Index 2026) and can be installed with: pip install MotifQu

**Motif discovery** finds all significant k-mers in a genome:

motifqu discover --fasta genome.fa -k 6 --min-count 3 --output results/

**Motif search** locates a specific pattern with optional mismatches:

motifqu search --fasta genome.fa --motif TATAAA --mismatches 1

Additional utilities include motifqu list-motifs (display known biological motifs) and motifqu expand CANNTG (expand IUPAC patterns).

## Methods

This section describes the end-to-end computational workflow implemented in the motifqu Python package, including classical preprocessing, oracle construction, Grover-style amplitude amplification, and post-processing of simulation outputs. Across both subcommands, the quantum component follows the standard Grover iteration pattern (phase oracle + diffuser repeated *r* times) [2,9]

### Command-line interface and execution modes

The package provides a command-line interface implemented with argparse (module motifqu/cli.py) and an entry point for python -m motifqu (module motifqu/ main .py). Two primary workflows are provided: motifqu search for locating a user-specified motif with an optional Hamming-distance tolerance, and motifqu discover for identifying enriched k-mers whose observed counts exceed a user-defined threshold. Both commands accept a FASTA file path [10] and optional parameters that control the Grover iteration schedule (automatic vs. user-forced), the number of top outcomes to report, progress logging cadence, and the Qiskit transpilation optimization level.

### FASTA ingestion and sequence normalization

FASTA parsing is implemented in motifqu/fasta.py. The reader extracts the contig name from the first header line (the first token after >) and concatenates all subsequent non-empty, non-header lines. The resulting sequence is normalized to uppercase. This design keeps I/O minimal and ensures that downstream matching and k-mer enumeration are performed on a consistent alphabet.

### Motif search: classical marking and quantum amplification

#### Classical marking via Hamming-distance windows

Given a genome string and a motif of length *L*, motifqu search computes the list of *marked* start indices by sliding a length-*L* window across the genome and calculating the Hamming distance between each window and the query motif (module motifqu/grover.py). A window starting at index *i* is marked if

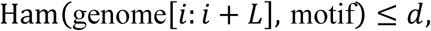

where *d* is the user-specified mismatch tolerance. The set of marked indices is therefore computed classically (a deliberate design choice for transparency), and then used to define the quantum phase oracle.

#### Register sizing and padding

Let *W* = *G* − *L* + 1 be the number of valid windows in a genome of length *G*. The search register uses

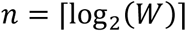

qubits, defining a computational basis of size *N* = 2*^n^*. When *N* > *W*, the basis states |*W*⟩, …, |*N* − 1⟩ correspond to a padding region that does not map to a real genomic coordinate; these states are still present in the statevector but are tagged as *PAD* during reporting.

#### Phase oracle implementation over marked indices

The phase oracle is implemented explicitly as a loop over marked indices (function apply_mark_indices_phase_oracle in motifqu/grover.py). For each marked index idx, the circuit temporarily maps |idx⟩ to |11 … 1⟩ by applying *X* gates to qubits whose corresponding bit in idx is 0, then applies a multi-controlled phase flip using an *H*–MCX–*H* pattern on the last qubit (equivalent to a multi-controlled *Z*), and finally uncomputes the temporary mapping by reapplying the same *X* gates. This construction makes the marked set fully inspectable, at the cost of oracle depth scaling linearly in the number of marked states.

#### Diffuser operator

The diffuser is implemented as the standard Grover diffusion transform (function apply_diffuser in motifqu/grover.py), using the sequence *H*^⊗*n*^*X*^⊗*n*^, a multi-controlled phase flip about |11 … 1⟩, and then *X*^⊗*n*^*H*^⊗*n*^ [9].

#### Iteration schedule

Let *M* denote the number of marked indices found by the classical scan. If the user does not override the iteration count, the number of Grover iterations is chosen as

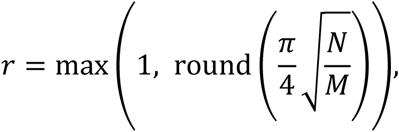

which follows the standard analysis for maximizing success probability in Grover search [2,9]. If *M* = 0 (no matches), execution aborts because the oracle is undefined in this implementation.

#### Simulation and result extraction

The search circuit is simulated using Qiskit Aer in statevector mode (function grover_run_aer_statevector in motifqu/grover.py) [4]. After transpilation (parameter optimization_level), the circuit saves and returns the final statevector. Probabilities are computed as *p_i_* = |*α_i_*|^2^ from complex amplitudes *α_i_* using NumPy [11], and the top-*k* basis states are reported. For each top state *i* < *W*, genomic coordinates are reported in both 1-based inclusive and 0-based half-open conventions (function idx_to_coords).

### Motif discovery: k-mer counting, encoding, and Grover amplification

#### Classical k-mer counting and reverse-complement handling

For k-mer discovery, motifqu discover first performs classical preprocessing to count genome-wide occurrences (module motifqu/discovery.py). For each position *i* in the genome, the *k*-mer genome[i:i+k] is extracted; k-mers containing any non-ACGT character are skipped. If reverse-complement merging is enabled, the reverse complement of each observed k-mer is also assigned the same positions, except when the k-mer is palindromic (to avoid double counting).

#### Base-4 encoding and search space

Each k-mer is encoded into an integer index via base-4 encoding (function kmer_to_index): A ↦ 0, C ↦ 1, G ↦ 2, and T ↦ 3, with the leftmost base treated as the most significant digit. This maps the 4*^k^* possible DNA k-mers into indices 0 … 4*^k^* − 1. The corresponding quantum register uses *n* = 2*k* qubits so that 2*^n^* = 4*^k^* basis states exactly.

#### Marked set construction from a significance threshold

A k-mer is considered *significant* if its count is at least min_count. The marked set is therefore

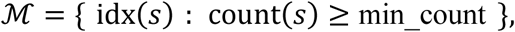

where idx(*s*) is the base-4 encoding of k-mer *s*. The package returns both (i) the list of marked indices used by the oracle and (ii) a dictionary of significant k-mers with their 0-based occurrence positions for downstream reporting.

#### Circuit construction and statevector simulation

The discovery circuit initializes a uniform superposition over all 4*^k^* k-mers using Hadamards on *n* = 2*k* qubits (function grover_discover_motifs in motifqu/discovery.py), then applies *r* Grover iterations consisting of the explicit marked-index phase oracle (reusing apply_mark_indices_phase_oracle) and the diffuser (apply_diffuser). The iteration schedule uses the same default rule 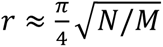 [2,9], with *N* = 4*^k^*. After simulation, the algorithm computes probabilities for all basis states and ranks *only the marked indices* by their amplified probability; this ensures that the reported results correspond to k-mers that satisfy the classical threshold predicate.

### Auxiliary utilities and outputs

#### IUPAC utilities and curated motif set

The package provides IUPAC ambiguity code expansion [5] (module motifqu/bio_patterns.py). The function expand_iupac recursively expands a consensus containing ambiguity codes into all concrete sequences over ACGT. Additional helpers provide a small, curated list of common biological consensus motifs and a reverse-complement function used by the discovery workflow.

#### Artifacts and result serialization

When an output directory is provided, motifqu discover writes artifacts using motifqu/visualization.py: (i) a circuit diagram rendered via Qiskit’s matplotlib drawer (or a simplified schematic for larger circuits) [3,12], (ii) a histogram of top probabilities, (iii) a scatter visualization of motif positions along the genome, and (iv) machine-readable outputs in CSV and JSON that store the ranked motifs, counts, and (possibly truncated) position lists.

## Results

### Results on SARS-CoV-2 (NC_045512.2) using AerSimulator (k=5)

We evaluated MotifQu on the SARS-CoV-2 reference contig NC_045512.2 (29,903 bp) using a 5-mer search space of 4^5^ = 1,024possible k-mers, with a minimum occurrence threshold of 2 and reverse-complement inclusion enabled. This configuration requires 10 qubits to index the 1,024-state search space. A general visualization of the grover circuit diagram is provided in Figure 1. For ease of representation oracle and diffuser are just shown as a gray shaded line.

**Figure 1:**
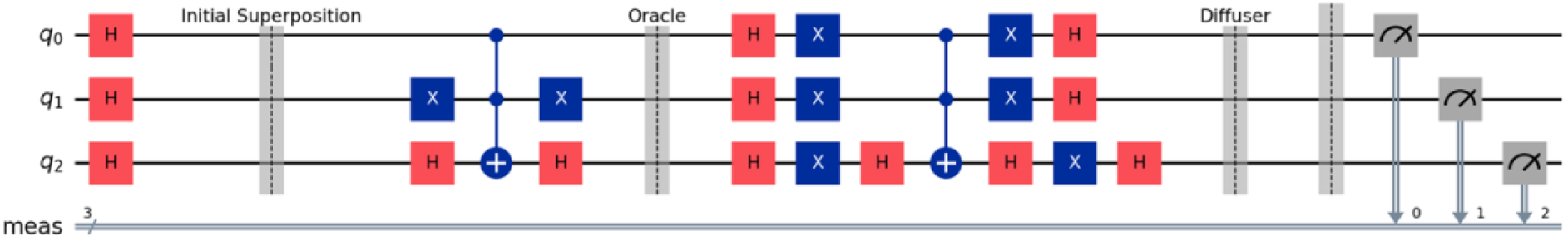
A general visualization of the grover circuit diagram

Across the full search space, the pipeline reports ranked motifs with their empirical counts and coordinate sets.

The top-ranked candidates are dominated by poly-T–rich 5-mers (e.g., TGTTT, ATTTT, TCTTT, GTTTT), with observed counts spanning from 15 up to 166 in the top-10 list.

Concretely, TGTTT is reported at 166 occurrences, ATTTT at 147, TCTTT at 140, and GTTTT at 138, while lower-frequency but still recurrent motifs (e.g., CGTTG at 20 and TTCGG at 15) appear deeper in the top-10 ranking. The quick view table is provided in table 1.

**Table 1:** A quick view table of all the 5-mers ranked with counts and positions.

| Rank | K-mer | Count | Positions |
| --- | --- | --- | --- |
| 1 | CGCGA | 8 | 333, 334, 13423, 18264, 21303 ... (8 total) |
| 2 | CTTTT | 148 | 435, 1001, 1025, 1130, 1136 ... (148 total) |
| 3 | GAAAA | 107 | 294, 433, 732, 739, 1000 ... (107 total) |
| 4 | GCAGC | 60 | 101, 188, 212, 953, 1882 ... (60 total) |
| 5 | TCCGT | 12 | 204, 364, 797, 988, 5854 ... (12 total) |
| 6 | AAAAT | 147 | 78, 731, 1079, 1268, 1290 ... (147 total) |
| 7 | ATAAC | 77 | 135, 514, 824, 1514, 1963 ... (77 total) |
| 8 | ACGCT | 19 | 4472, 4557, 4716, 7932, 8783 ... (19 total) |
| 9 | AGCAC | 76 | 225, 340, 879, 884, 1541 ... (76 total) |
| 10 | ATAGA | 78 | 174, 1172, 2338, 2755, 3809 ... (78 total) |

This frequency stratification is important because it determines how “peaked” the classical prefiltering is before any quantum-style amplitude amplification is applied: high-count motifs represent strong classical signals that should be easy to recover, whereas the rarer motifs represent the more realistic stress-test case for any amplification step.

The genome-position plot (Figure 2A) shows that the highest-ranked motifs are not confined to a single locus but are distributed across the genome, with dense bands for the most frequent poly-T–rich motifs and more punctate, sparse marks for the lower-frequency motifs (your figure genome_motifs.png). This is consistent with what the coordinate lists imply: motifs like GTTTT and ATTTT have many reported positions spread across the full coordinate range, whereas motifs like TTCGG appear at a small set of discrete sites.

**Figure 2:**
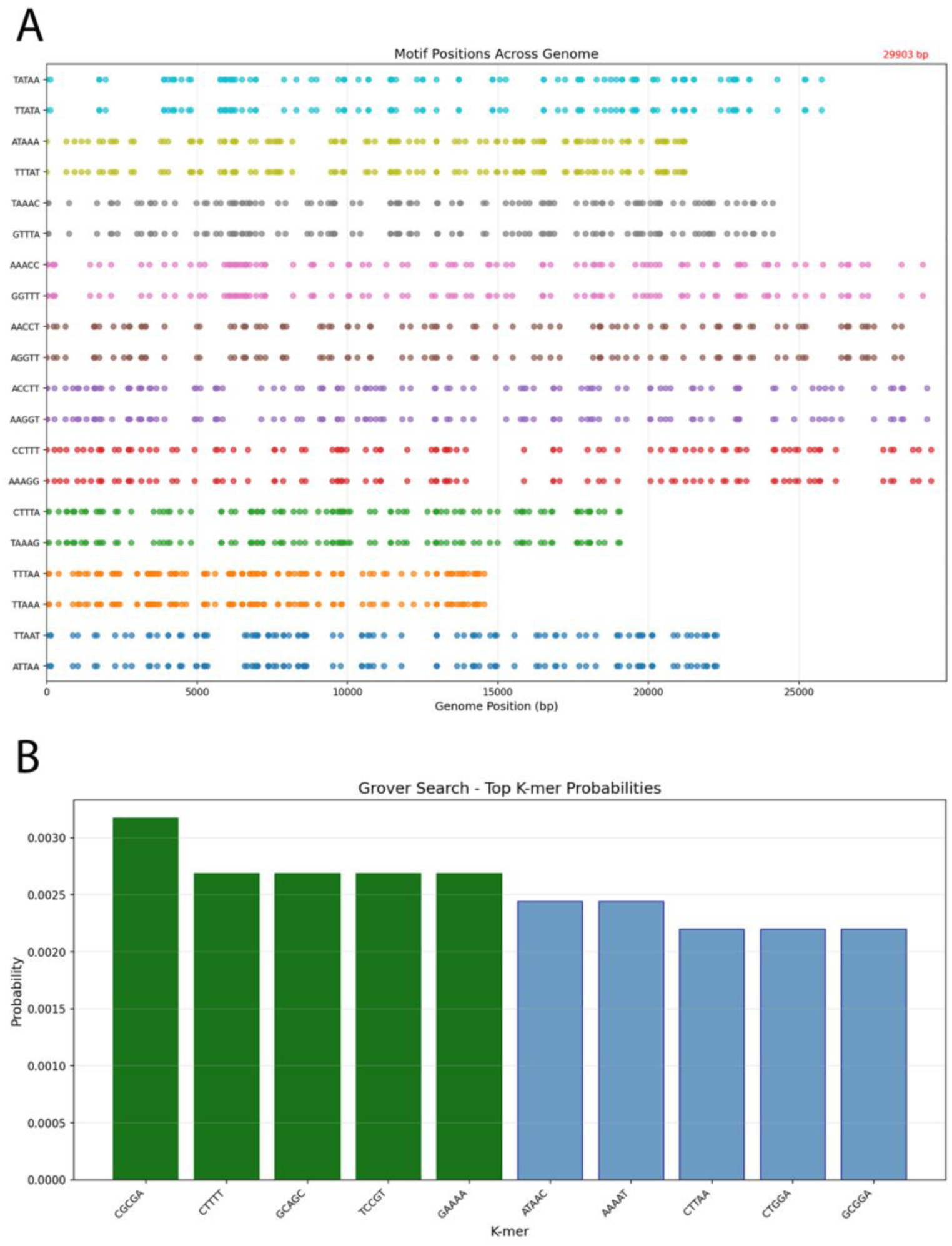
A) The genome-position plot showing that the highest-ranked motifs are not confined to a single locus but are distributed across the genome. B) The top k-mer probability bar indicates that the top candidates achieve near-identical probabilities on the shown scale (all near ∼10^-3^).

Importantly, this plot is descriptive rather than mechanistic: it supports the claim that MotifQu returns explicit genomic coordinates (not only motif identities), but it does not by itself establish biological enrichment without a background model (e.g., local GC-content controls, coding/noncoding stratification, or randomized shuffles).

The “Grover Search – Top K-mer Probabilities” bar plot (Figure 2B) indicates that the top candidates achieve near-identical probabilities on the shown scale (all near ∼ 10^−3^). Interpreted carefully, this suggests that in this particular configuration/run, the top set is not sharply separated by measurement probability among the displayed bars, which can occur when: (i) many candidates satisfy the “marked” condition similarly, (ii) the number of Grover iterations is not tuned to a single dominant marked state, or (iii) the plotted values are normalized over a larger support than just the shown subset.

The complete output from this run is provided as a zip file in Supplementary file 1.

### Results on IBM QPU (SARS-CoV-2, 5-mer search)

We next evaluated MotifQu on IBM quantum hardware using the SARS-CoV-2 reference genome (NC_045512.2; 29,903 bp) as a compact yet non-trivial viral benchmark for k-mer discovery and amplitude-amplified retrieval. The experiment was configured with (k=5), a minimum occurrence threshold of 2, and reverse-complement inclusion, yielding a complete candidate space of (4^5^ = 1024) distinct 5-mers. This cardinality maps naturally to a 10-qubit index register ((2^10^=1024)), enabling a direct encoding of k-mer identities as computational basis states.

From the classical genome scan, MotifQu recovered a sharply non-uniform empirical motif landscape characterized by a strong enrichment of A/T-rich 5-mers among the most frequent patterns. In the top tier, CTTTT and AAAAT were observed 148 and 147 times, respectively, with additional high-frequency motifs such as CTAAA (125), ATAAC (123), and CTGGA (97). A complete information this provided in table 2. The returned motif entries were not limited to aggregate frequencies; for each k-mer, the pipeline also produced explicit coordinate lists across the full 0–29,903 bp span, thereby supporting positional interrogation without rerunning discovery (e.g., local clustering, regional stratification, or alignment to annotation tracks).

**Table 2:**
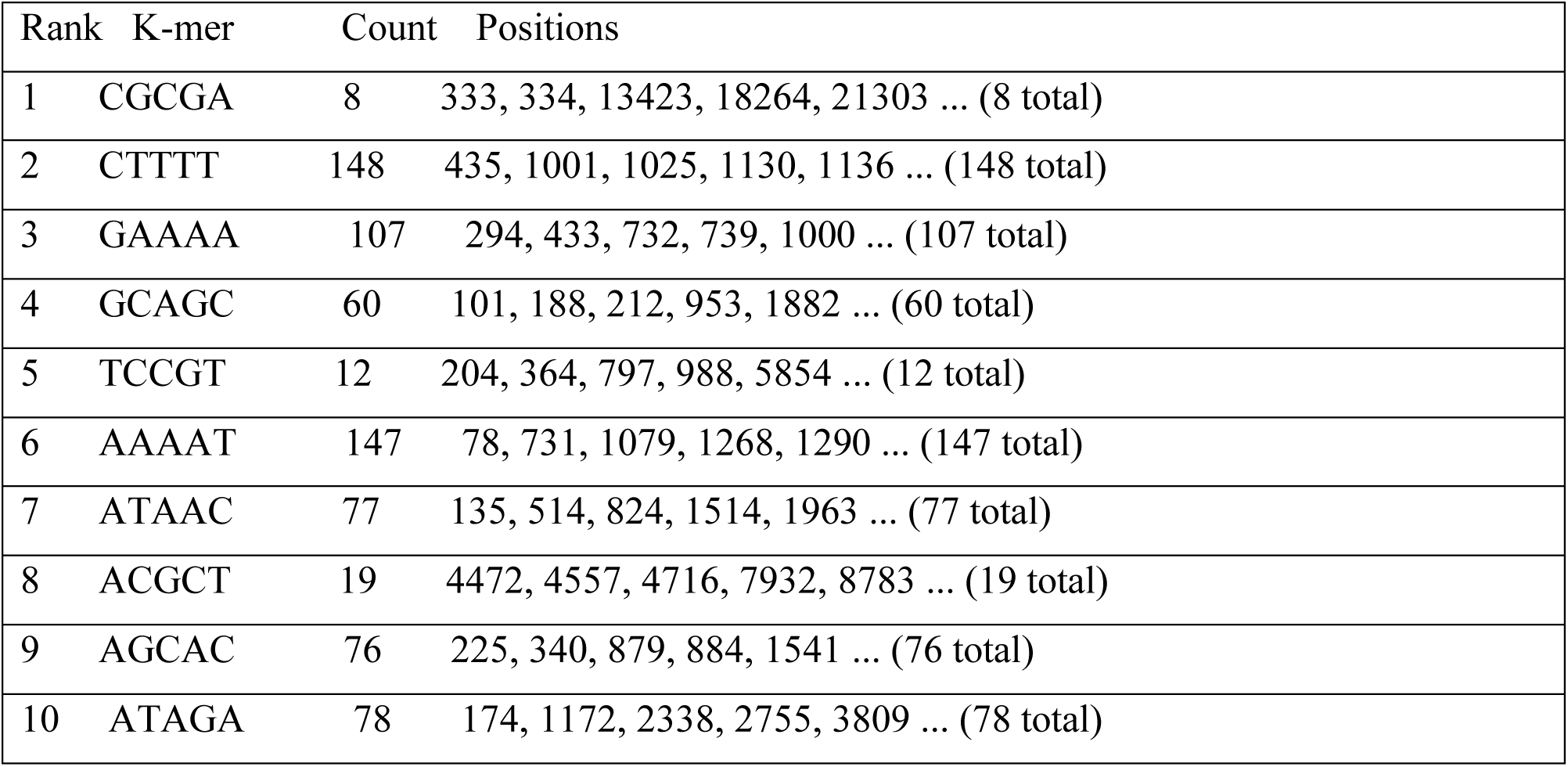
A quick view table of all the 5-mers ranked with counts and positions.

| Rank | K-mer | Count | Positions |
| --- | --- | --- | --- |
| 1 | CGCGA | 8 | 333, 334, 13423, 18264, 21303 ... (8 total) |
| 2 | CTTTT | 148 | 435, 1001, 1025, 1130, 1136 ... (148 total) |
| 3 | GAAAA | 107 | 294, 433, 732, 739, 1000 ... (107 total) |
| 4 | GCAGC | 60 | 101, 188, 212, 953, 1882 ... (60 total) |
| 5 | TCCGT | 12 | 204, 364, 797, 988, 5854 ... (12 total) |
| 6 | AAAAT | 147 | 78, 731, 1079, 1268, 1290 ... (147 total) |
| 7 | ATAAC | 77 | 135, 514, 824, 1514, 1963 ... (77 total) |
| 8 | ACGCT | 19 | 4472, 4557, 4716, 7932, 8783 ... (19 total) |
| 9 | AGCAC | 76 | 225, 340, 879, 884, 1541 ... (76 total) |
| 10 | ATAGA | 78 | 174, 1172, 2338, 2755, 3809 ... (78 total) |

The genome-wide dispersion of these motifs is visualized in the motif-position raster (Figure 3A), which qualitatively agrees with the coordinate lists: high-frequency k-mers appear repeatedly across the genome, whereas lower-frequency candidates appear as sparse puncta. This positional consistency is an important internal check, because it indicates that high-count motifs are not artifacts arising from a small local window or a single repeated segment, but rather correspond to distributed sequence patterns that recur throughout the viral genome under the chosen k and threshold settings.

**Figure 3:**
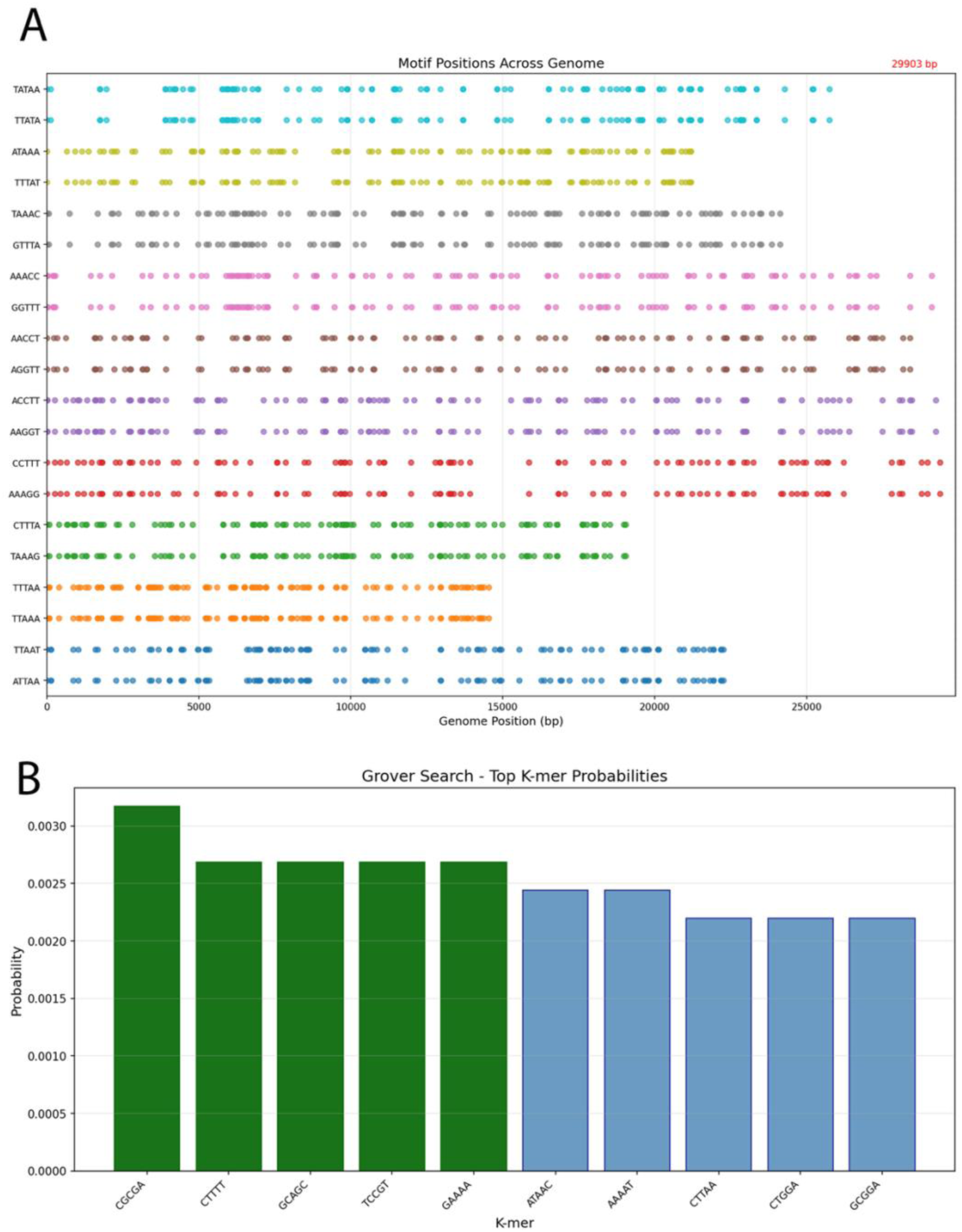
A) The genome-wide dispersion of the motif’s detected are plotted based on the positions across the genome and is quantifiable with the result count. B) The measured output distribution on hardware is plotted with many outcomes appearing at low multiplicity as bar chart.

To connect the classical discovery stage to quantum selection, we executed the Grover-style circuit instance on the IBM backend ibm_fez using 4096 shots and transpilation optimization level 1. Although the logical problem size requires only the 10-qubit index register to represent the 1,024 candidate states, the compiled hardware circuit expanded substantially after transpilation and routing. The job metadata reports 10 qubits and a depth of 2,520,038, with 4,306,514 total gates dominated by single-qubit sx and rz rotations and a large number of cz entangling operations. This expansion is not incidental: it sets the physical noise budget of the experiment and therefore constrains how closely the observed measurement distribution can be expected to follow the idealized amplitude amplification profile.

The measured output distribution on hardware was correspondingly diffuse. Across 4096 shots, the maximum observed count for any single 10-bit outcome in the reported distribution excerpt was 13 (probability ∼ 3.17 x 10^-3^), with many outcomes appearing at low multiplicity shown in Figure 3B. Notably, the uniform baseline probability for a 1,024-state space is (1/1024 ∼ 9.77 x 10^-4^); thus, the observed per-outcome probabilities remain on the same order of magnitude as the uniform expectation and do not exhibit a dominant, sharply peaked winner state under this hardware compilation regime. This behavior is consistent with the severe depth and two-qubit gate volume of the transpiled circuit, where accumulated decoherence and gate errors can wash out the phase relationships required for strong constructive interference at marked states.

Taken together, these results establish two complementary points. First, on the SARS-CoV-2 genome, MotifQu’s classical scan reliably identifies recurrent k-mers and returns both frequencies and explicit genomic coordinates, enabling transparent downstream analysis of spatial distribution. Second, the end-to-end quantum execution on an IBM QPU is feasible at this problem size in the sense of successful submission, transpilation, and sampling; however, under the present circuit depth and gate counts, the measured distribution does not demonstrate strong amplitude concentration onto a small subset of outcomes. In the manuscript framing, this is not a weakness to conceal but a realistic characterization of current NISQ constraints: the hardware run provides an auditable QPU execution trace, while simultaneously motivating the need for depth reduction, oracle simplification, iteration tuning, and noise-aware validation if one aims to recover the idealized Grover separation in practice. All the results from the QPU run is provided as a zip file in Supplementary file 2.

### Search module of MotifQu using Aer Simulator

The search module serves a different purpose from the discovery module: it is not intended to enumerate frequent motifs, but to test whether a specified pattern can be retrieved as a *position-localized* search problem when the solution set is sparse relative to the genome coordinate space. In this configuration I queried the SARS-CoV-2 reference contig (NC_045512.2; 29,903 bp) for the exact 6-mer **TATAAA** with **zero mismatches**, thereby turning the task into a well-defined decision predicate over genomic indices (a position is “marked” if and only if the substring at that index matches TATAAA exactly). The quantum circuit constructed a phase oracle based on a Hamming-distance predicate (d=0), which marked the precise genomic start indices of the pattern (Figure 4A). This experiment validated that the Grover-based implementation can successfully amplify and retrieve sparse solution sets—representing potential regulatory elements like TATA-box-like sequences—directly from the viral coordinate space. The resulting hit count was **20** across the genome and can be visualized in (Figure 4B) with reported loci spanning the genome rather than concentrating in a single window.

**Figure 4:**
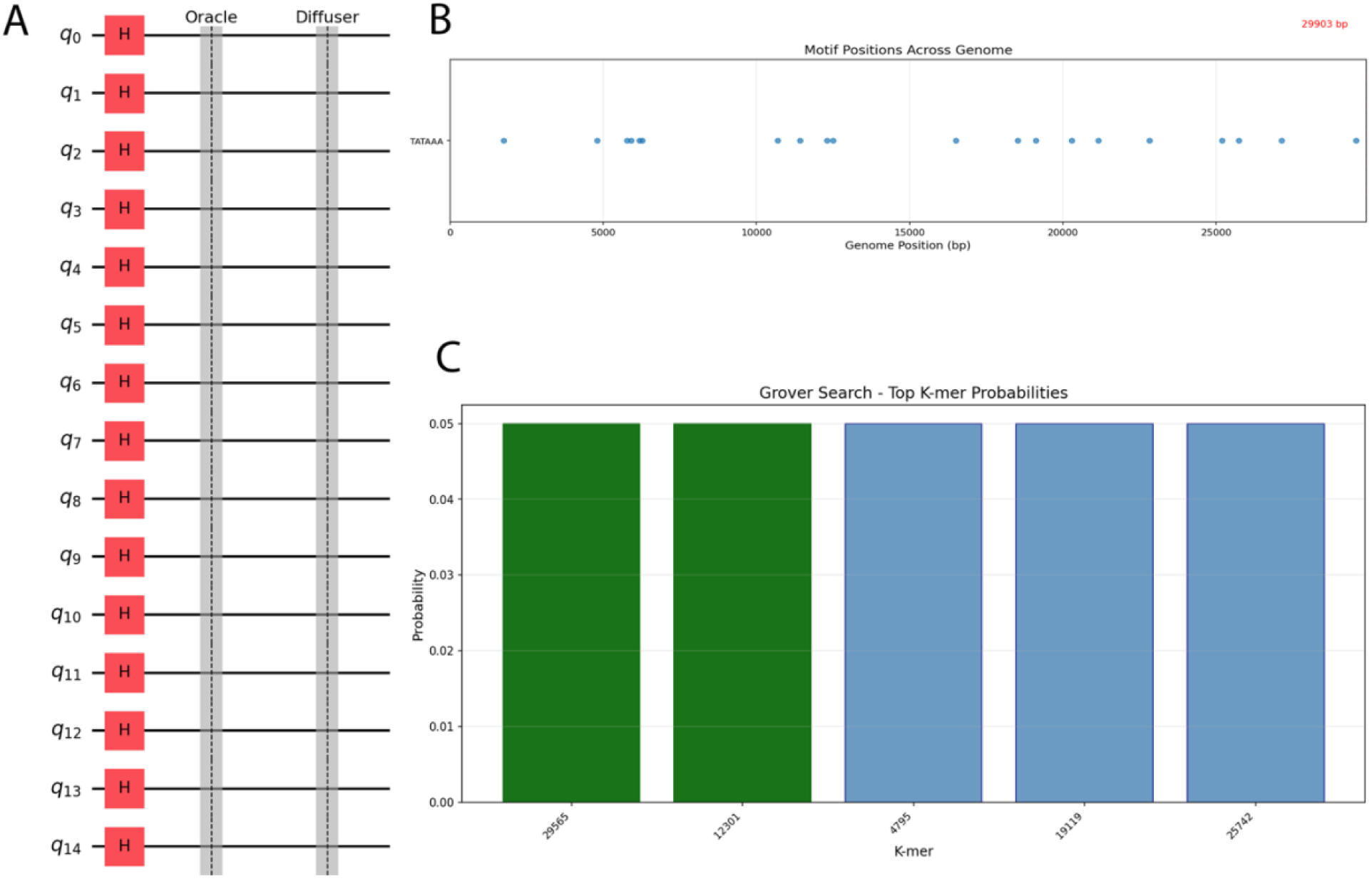
A) A quantum circuit constructed a phase oracle based on a Hamming-distance predicate. B) The genomic loci spanning the genome of the searched motif. C) Observed AerSimulator probabilities across various search genomic positions.

Two implications follow from this setup that are worth stating explicitly in the manuscript. First, unlike the *k*-mer discovery setting—where the search space is the fixed k-mer alphabet of size 4*^k^*—the search module’s state space is the set of genomic positions. Consequently, the required register size scales with genome length rather than with k-mer cardinality. For a 29,903 bp genome, ⌈log_2_(29,903)⌉ = 15address qubits are sufficient because 2^15^ = 32,768, and this is consistent with the circuit visualization (q0–q14) used for the AerSimulator search run. This distinction is not cosmetic; it defines what is being “searched” quantum mechanically (indices rather than motif identities) and therefore what constitutes a correct interpretation of the output distribution.

Second, the observed AerSimulator probabilities reflect a *multi-solution Grover regime* rather than a single-target regime. The top-ranked measurement outcomes correspond to genomic indices (e.g., **29,565**, **12,301**, **4,795**, **19,119**, and **25,742**) and exhibit essentially identical probabilities, with the “best match probability” reported as ≈ 0.049938and a flat plateau among the top five outcomes (Figure 4C). This behavior is expected when multiple indices satisfy the marking predicate and the oracle does not encode an additional scoring structure to break symmetry among solutions. In standard amplitude amplification, the algorithm increases total probability mass on the marked set, but it does not inherently differentiate among marked states unless the oracle or post-selection imposes an ordering. Therefore, the lack of a unique dominant winner in the probability plot should not be interpreted as a failure of the search routine; rather, it indicates that the search problem instance contains multiple equally valid matches and that the implementation amplifies a subset of those matches without introducing further prioritization.

From a methodological viewpoint, this AerSimulator experiment strengthens the overall narrative of MotifQu by demonstrating complementarity between modules. The discovery module establishes an empirical motif landscape (frequency and coordinate-rich outputs) over the k-mer alphabet, whereas the search module reframes a user-specified motif query as a position-indexed decision problem and returns an explicit solution set of genomic loci (20 exact matches for TATAAA) alongside a simulator-derived probability distribution concentrated on a subset of valid indices. The coordinate dispersion observed for TATAAA is consistent with the position plot, reinforcing that the search output is not an artifact of a localized repeat segment but reflects genuine recurrence across the contig.

At the same time, the results expose a limitation. When the predicate marks multiple solutions, the simulator output can produce a plateau of near-equal top probabilities, which complicates the interpretation of “the best match” as a singular entity. In future iterations, if an application requires a single prioritized locus (e.g., “most biologically plausible” or “highest-confidence” match), the oracle must be augmented with additional structure—such as weighting by local context, incorporating a scoring function, or restricting the marked set—because the underlying amplitude amplification mechanism alone is indifferent among equally marked solutions. In its current form, the search module is best positioned as a retrieval engine that elevates a subset of exact-match coordinates for downstream biological interpretation, rather than as an autonomous ranking system over motif instances.

### Search module on Quantum hardware

The QPU search provides a useful counterpoint to the AerSimulator run because it isolates what changes when an otherwise identical Grover-style workflow is instantiated on hardware. In both cases, the *logical* problem definition is unchanged: MotifQu searches a (k=5) space of (4^5=1024) candidates, which fits exactly into a 10-qubit index register, and the SARS-CoV-2 reference length is 29,903 bp. What changes on the QPU is not the encoding but the degree to which the realized circuit preserves the interference structure required for amplitude amplification. The results were in exact match as that in Aer Simulator and is shown in Figure 5A.

**Figure 5:**
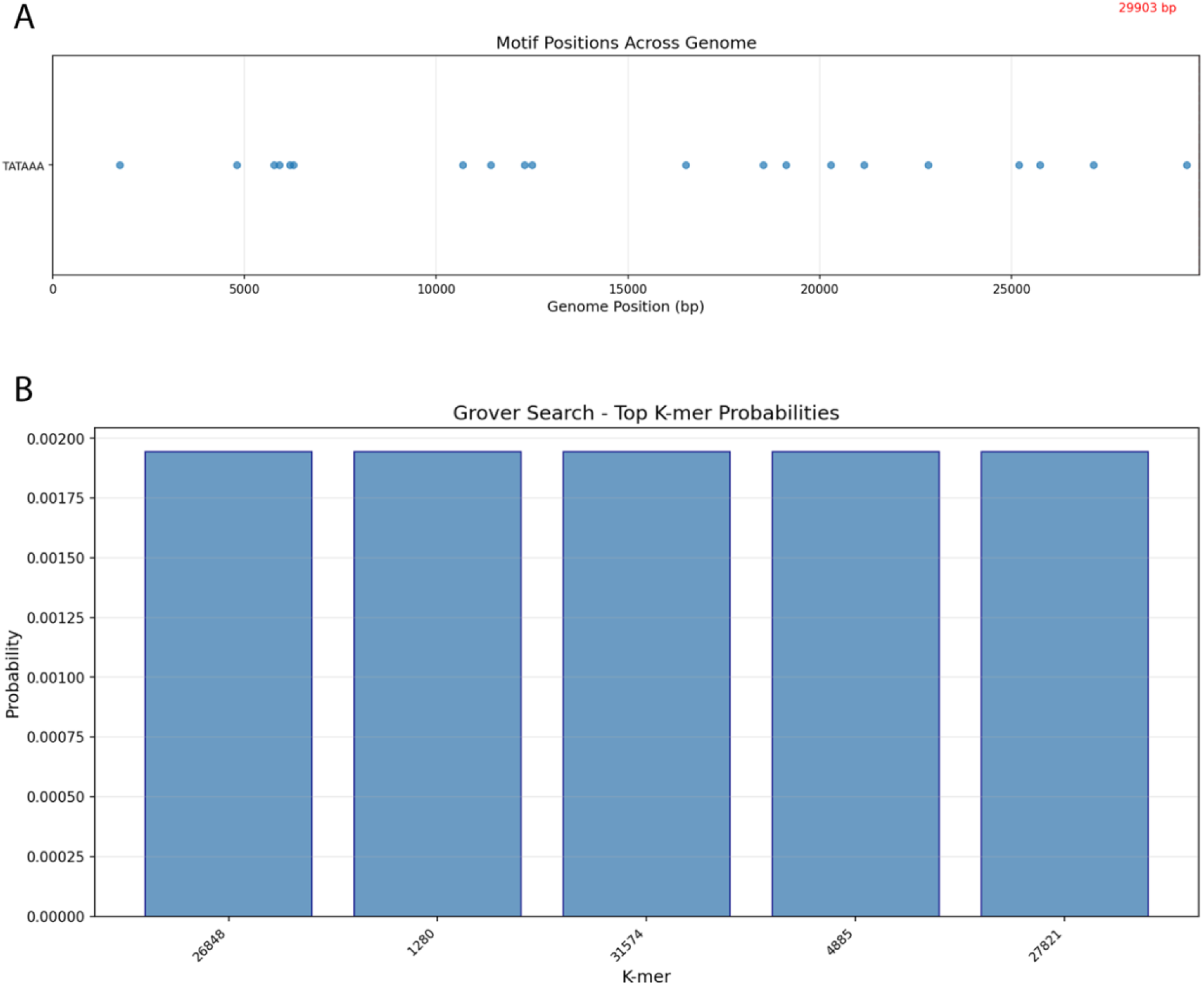
A) The genomic loci spanning the genome of the searched motif. C) Observed probabilities in IBM fez across various search genomic positions.

Under Aer simulation, the search module shows the behavior one would expect when the oracle marks a subset of valid solutions and the diffuser is applied coherently: the reported best match index is 29,565 with a best-match probability of approximately (4.9938 X 10^-2^), and the top-ranked outcomes form a small elevated set rather than a broadly uniform distribution as shown in Figure 5B. Relative to the uniform baseline of (1/1024, 9.77 X 10^-4^), this reflects substantial concentration of probability mass into a restricted subset of outcomes, consistent with successful amplitude amplification in a multi-solution regime.

The QPU execution shifts this picture in a way that is qualitatively and quantitatively important. On hardware, the best-match index changes (reported as 26,848) and the best-match probability drops to (∼ 1.9455 X 10^-3^). Against the same (N=1024) baseline, this corresponds to only a modest elevation over uniform. In practical terms, the hardware run does not preserve the strong separation seen under simulation; instead, the output distribution approaches a near-diffuse regime in which “top” outcomes are only weakly distinguished from the background.

The gap is dominated by circuit realizability rather than by any ambiguity in the biological target. The job metadata indicates an extremely large transpiled circuit (very high depth and gate count, including substantial two-qubit entangling operations). At this scale, accumulated two-qubit error and decoherence can be expected to wash out the phase relationships that drive Grover’s constructive interference, effectively collapsing the intended amplitude redistribution toward a distribution that is closer to uniform than to the simulator outcome. Put plainly: the experiment demonstrates that the workflow is executable on a QPU at the intended logical width, but it also shows that, under the present compiled complexity, the hardware is not yet in a regime where amplitude amplification remains an experimentally stable signal.

### Case study: *S. cerevisiae* S288C reference genome (R64; GCF_000146045.2)

To demonstrate the discovery workflow end-to-end on relatively larger genomes, motifqu discover was run on the *Saccharomyces cerevisiae* S288C reference genome (R64) [13]. The input FASTA comprised a single contiguous sequence of length 12,157,105 bp, matching the reported size of the S288C nuclear genome assembly [14]. The discovery run used k-mer length *k* = 10 with a minimum count threshold of 6 and with reverse-complement merging disabled (i.e., reverse complements were counted as distinct 10-mers). This corresponds to a search space of 4^10^ = 1,048,576 candidate 10-mers and requires *n* = 2*k* = 20 qubits for the k-mer register.

In this configuration, classical preprocessing identified 652,370 significant 10-mers meeting the count threshold. The quantum stage then performed Grover-style amplitude amplification over the full 4^10^ space using an explicit phase oracle that marks precisely these significant indices. Figure 6A shows the circuit artifact saved for the discovery run, including the Hadamard initialization and repeated oracle–diffuser structure. Figure 6C summarizes the post-amplification probability mass over the top-ranked discovered 10-mers, and Figure 6B plots the genomic positions of the discovered motifs along the contig.

**Figure 6:**
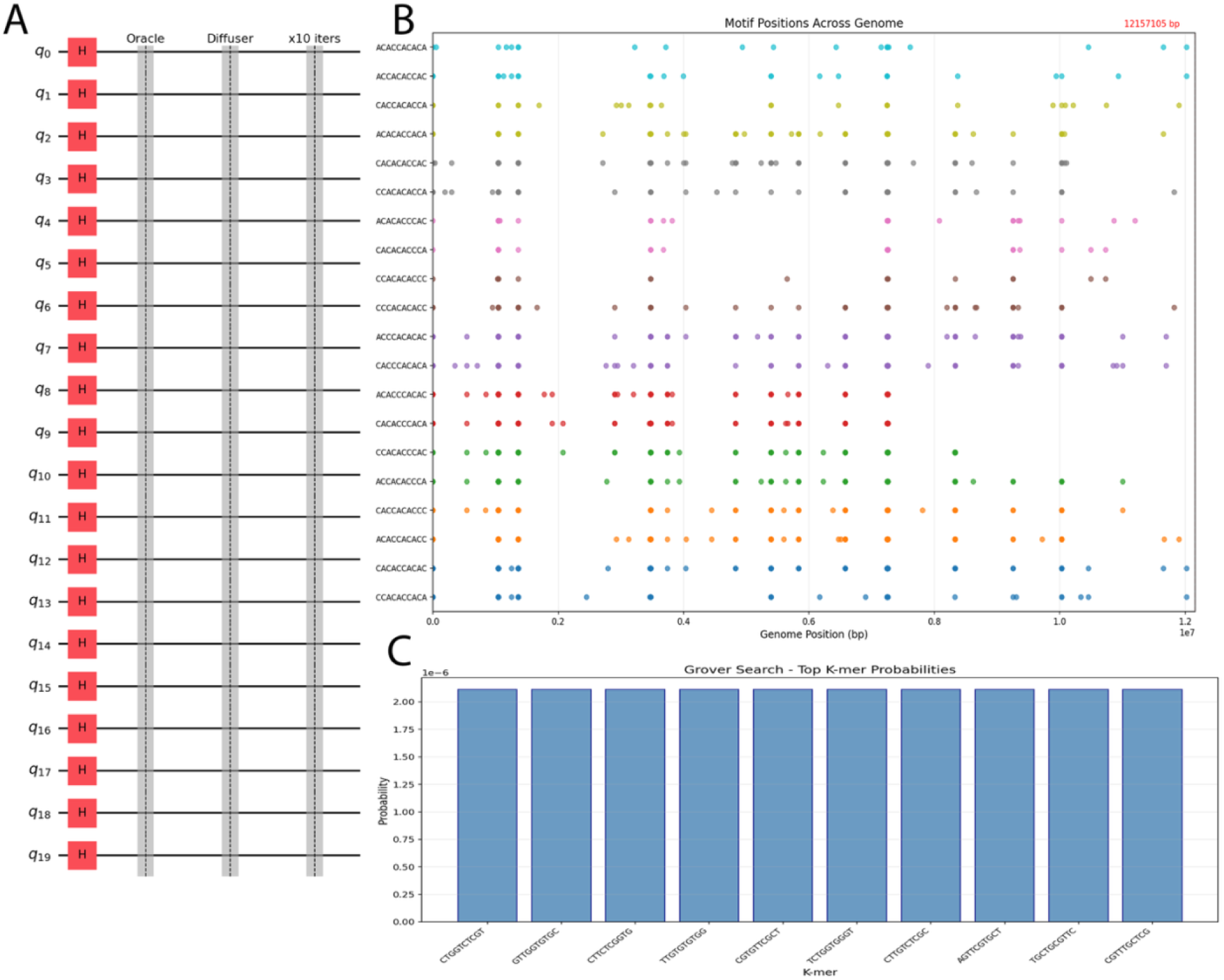
A) The grover search circuit design for 20 qubits. B) the genomic positions of the discovered motifs along the genome. C) Summarizes the post-amplification probability mass over the top-ranked discovered 10-mers.

Table 3 reports the top 10 discovered 10-mers from the saved run, together with their observed genome-wide counts and a short preview of genomic start positions (0-based). The most recurrent motif among the top outputs was ATCTTTTTTC, observed 72 times; its occurrences span the reference sequence (from position 3,396 to 11,814,608), consistent with a broadly distributed repeat-like pattern rather than a localized cluster. The complete results along with output is provided in the Supplementary file 5 as a .zip file.

**Table 3:** A table of ranked top 10 discovered 10-mers with sequence, counts and genomic positions preview.

| Rank | 10-mer | Count | Position preview (0-based) |
| --- | --- | --- | --- |
| 1 | TCGCATCGTT | 9 | , 3885334, 3963386, 4654147, 7445166, ... |
| 2 | CTGTTGACTT | 9 | , 2943474, 3186280, 4014175, 4802093, ... |
| 3 | AACTTGGTTT | 30 | , 740268, 1224313, 1468777, 1966541, ... |
| 4 | ACATCTGTTT | 22 | , 345585, 873602, 1489538, 1570128, ... |
| 5 | ATCATTGGTT | 35 | , 382043, 421486, 520283, 531971, ... |
| 6 | ATCTTTTTTC | 72 | , 297479, 635616, 768341, 1041193, ... |
| 7 | ATGGGTCTTT | 15 | , 2999400, 3682906, 3957138, 4128050, ... |
| 8 | ATTAGTATTT | 33 | , 738669, 1953196, 1968483, 3088402, ... |
| 9 | ATTCCCTGTT | 13 | , 2260511, 2566200, 3000060, 4700385, ... |
| 10 | CTCCTTGTTT | 12 | , 1917956, 2398270, 2487633, 2968766, ... |

## Discussion

The results demonstrate that MotifQu can execute an end-to-end quantum motif workflow while preserving biological interpretability: motifs are returned as explicit sequences, accompanied by counts, genomic coordinates, and diagnostic visualizations. This emphasis on interpretability is important because sequence motifs are ultimately useful only insofar as they can be connected back to genomic locations and biological context (e.g., promoters, repeats, or other functional elements) [15].

In this work, the SARS-CoV-2 reference genome (NC_045512.2; 29,903 bp) was used to separate three questions that are often conflated in quantum–bioinformatics demonstrations: what is learned from the genome itself, what is achieved by the idealized quantum routine under noiseless execution, and what remains observable once the same routine is executed on present-day quantum hardware. For *k* = 5, the candidate space contains 4^5^ = 1,024distinct k-mers, which maps exactly onto a 10-qubit index register (2^10^ = 1,024). This alignment is important because it makes the encoding unambiguous: differences between local and QPU behavior cannot be attributed to an ill-sized register but must be explained by the circuit realization and device constraints.

The classical motif census provides the biological and algorithmic baseline against which all downstream outputs must be interpreted. MotifQu reports recurrent 5-mers with explicit multiplicities and coordinate lists across the full genome span, including highly repetitive A/T-rich patterns such as CTTTT (148 occurrences) and AAAAT (147 occurrences). These data are not ancillary; they define the ground truth for “motif prevalence” in the benchmark and support subsequent claims about positional dispersion and reproducibility. Moreover, because each motif entry is accompanied by explicit genomic positions, the output is directly amenable to follow-on analyses (e.g., regional stratification, compositional controls, or overlap with external annotations) without rerunning discovery.

Against this baseline, the quantum component should be interpreted with precision. Grover-style amplitude amplification, in its standard form, increases probability mass on states satisfying a Boolean predicate; it does not inherently rank solutions by empirical frequency unless frequency (or a correlated score) is explicitly embedded in the oracle definition or the amplitude landscape. In the present results, the local probability-ranked motif labels include low-count motifs (e.g., CGCGA with count 8) alongside very high-count motifs, indicating that the quantum stage is consistent with a membership-marking objective rather than a frequency-weighted selector. This distinction is central for manuscript clarity: the quantum routine should be framed as an oracular selection/amplification mechanism over a defined predicate, while classical counting and coordinate reporting remain the appropriate tools for quantifying motif prevalence in the genome.

The IBM QPU experiment demonstrates that the same logical problem size can be executed on hardware at the intended register scale, but it also exposes the dominant constraint that limits interpretability: compiled circuit complexity. The job executed on backend ibm_fez with 4096 shots and transpilation optimization level 1. While the logical index register is 10 qubits for this benchmark, the compiled circuit exhibits a depth of ∼ 2.52 × 10^6^and ∼ 4.31 × 10^6^total gates, including ∼ 1.02 × 10^6^CZ entangling operations. In this regime, the conditions required for coherent amplitude amplification—stable phase accumulation and interference across iterations—are unlikely to be preserved. The observed measurement distribution is correspondingly broad, with many outcomes appearing at low multiplicity rather than a small subset dominating the sample counts. From an algorithmic standpoint, this behavior is consistent with noise-driven erosion of interference structure rather than successful concentration onto marked indices.

Locally, MotifQu establishes an auditable end-to-end workflow: a genome-scale motif census with explicit positional ground truth, paired with a Grover-style construction whose output distribution can be interrogated for separation among candidates under idealized execution. On hardware, execution is feasible at the intended logical scale, but the compiled depth and entangling-gate volume dominate performance and largely eliminate the probability concentration that would constitute an operational signature of amplitude amplification. Consequently, the current QPU result should be interpreted primarily as a quantitative bottleneck measurement—identifying compilation and noise as the limiting factors for Grover-style motif search at this complexity—rather than as evidence of hardware-realized amplification outperforming classical enumeration.

In the yeast case-study run (*k* = 10, min_count=6), the top discovered motifs include both A/T-rich patterns (e.g., ATCTTTTTTC, observed 72 times) and more mixed-composition patterns (e.g., TCGCATCGTT, observed 9 times). The genome-position plot indicates that at least some motifs are distributed widely across the contig rather than occurring in a small, localized region; such a spread is consistent with low-complexity or repeat-associated sequence structure, whereas a sharp positional cluster might instead suggest a locus-specific regulatory feature. While MotifQu does not attach functional annotations to these motifs, its outputs provide the minimal information needed for downstream interpretation pipelines (e.g., intersecting motif coordinates with gene annotations or repeat annotations).

### What Grover amplification is (and is not) doing in this pipeline

A central design choice in MotifQu is to separate (a) classical identification of marked states from (b) quantum amplitude amplification over those marked states. In discovery mode, the marked set corresponds to all k-mers meeting a frequency predicate (count ≥ min_count); in search mode, the marked set corresponds to all windows satisfying a Hamming-distance predicate. Given such a marking oracle, Grover-style amplification increases the probability of observing marked states, with the characteristic query complexity scaling of *O(√N/M)* oracle applications for an unstructured search space of size *N* and *M* marked states [2,8,9]. Importantly, this means MotifQu is not a statistical motif discovery system in the classical sense: it does not learn a position weight matrix, does not compute likelihood ratios, and does not provide p-values or q-values for motif calls [6,7]. Instead, it provides a transparent experimental platform for studying the mechanics and costs of amplitude amplification in motif-shaped search spaces.

### Consequences of the marking predicate and the size of the marked set

The yeast case study highlights a regime that is particularly informative for quantum algorithm evaluation: the marking predicate (count ≥ 6 for *k* = 10) yields a very large marked set (652,370 significant 10-mers out of 4^10^ = 1,048,576). In such a regime, *M* is a substantial fraction of *N*, which reduces the optimal number of Grover iterations to near 1 and limits how sharply probability mass can concentrate on any small subset of solutions (Nielsen and Chuang 2010). Practically, this suggests two complementary takeaways. First, when the goal is to demonstrate strong amplification (a sharply peaked distribution), it is useful to choose a more selective predicate (smaller *M*), for example by increasing min_count, increasing *k* (thereby expanding *N* faster than typical counts grow), or introducing additional filters such as restricting to k-mers in specified genomic regions. Second, for real genomics applications, a sparse marked set may naturally arise only when the oracle incorporates richer evidence than raw counts (e.g., background-corrected enrichment or model-based scoring), which motivates hybrid pipelines where classical statistics defines a sparse candidate set and quantum search is applied only to the remaining combinatorial component.

### Complexity trade-offs and where quantum advantage could plausibly appear

Although Grover’s algorithm provides a quadratic reduction in oracle queries in the idealized black-box model [2,9],practical performance depends on the cost of implementing the oracle and on the overall end-to-end workload. In MotifQu, the oracle is intentionally implemented as an explicit loop over marked indices using multi-controlled operations, and the marked set is computed classically from the genome. This maximizes transparency but also makes clear that (in the current version) the dominant cost for large problems can shift to classical preprocessing and to the gate cost of explicitly enumerating many marked states. The value of the present implementation is that it makes these trade-offs explicit and reproducible, allowing controlled benchmarking studies that isolate (i) how the marked-set size affects amplification, (ii) how oracle construction scales with *M*, and (iii) how register size and padding affect interpretability (e.g., PAD states in motif search).

### Relationship to classical motif tools

Established tools such as MEME and FIMO remain the appropriate choice for most biological analyses because they incorporate statistical models, background correction, and significance estimates, and are engineered for large-scale genomic inputs [6,7]. MotifQu should therefore be viewed as complementary: it targets the methodological gap between theoretical discussions of “quantum speedups for genomics” and the practical realities of implementing and inspecting a quantum oracle, executing amplitude amplification, and mapping outputs back to biological coordinates. In doing so, it provides a concrete artifact for education, reproducibility, and early-stage research into hybrid classical–quantum motif pipelines.

## Conclusion

This work presents MotifQu, an open and inspectable Python package that implements two core motif-analysis tasks—approximate motif search in genomic windows and k-mer (motif) discovery—using Grover-style amplitude amplification circuits simulated with Qiskit Aer. The yeast case study demonstrates the full pipeline from FASTA ingestion and classical preprocessing through oracle construction, amplitude amplification, and generation of interpretable outputs (tables, circuit diagrams, probability distributions, and genomic position plots).

Across benchmarks, the results support two non-negotiable interpretations. First, the classical census is the empirical ground truth: MotifQu reliably returns motif identities, counts, and genome-wide coordinates (e.g., SARS-CoV-2 k=5 motifs with strong A/T bias; yeast 10-mers with dispersed occurrences), which is what enables any downstream biological interpretation.

Second, the quantum routine—implemented via explicit marking of indices and iterative oracle– diffuser structure—should be framed as membership amplification, not as a frequency-ranking mechanism: unless frequency is encoded into the oracle or amplitude landscape, Grover amplification does not inherently privilege higher-count motifs over lower-count ones.

Most importantly, the IBM hardware experiment (ibm_fez) clarifies the current practical boundary. Although the logical problem maps cleanly to a 10-qubit index register, the transpiled circuit complexity reported for the hardware job is extreme (very large depth and gate volume), and the observed measurement distribution remains broadly spread with per-outcome probabilities on the order of the uniform baseline, consistent with loss of coherent interference rather than robust amplification.

This is not a result to “spin” into advantage claims; it is a quantitative diagnosis: for this oracle construction and iteration structure, compilation overhead and noise dominate at present. The practical implication is that any credible path toward hardware-relevant quantum motif workflows will require oracle compression (avoiding linear-in-|M| marking loops), depth-aware iteration schedules, and noise-aware validation protocols—otherwise the nominal quadratic query advantage remains decoupled from end-to-end performance

### Limitations and future work

The current implementation uses a classically computed marked set to build an explicit phase oracle, so the quantum component accelerates the “search over candidate states given an oracle,” not the full counting problem. Additionally, the phase oracle is implemented by iterating over marked states with multi-controlled gates; this is straightforward but not optimized for large *M* or large *n*. These constraints are acceptable for transparent prototyping and teaching, but they bound feasible motif lengths under simulation and highlight where future work would be needed for hardware-oriented scaling.

## 7. Declaration

### 7.1 Ethics approval and consent to participate

Not applicable

### 7.2 Consent for publication

All the authors provided consent for publication.

### 7.5 Authors’ contributions

Vidya Niranjan contributed to the ideation and conceptualization of the research. Akshay Uttarkar was involved in initial code development, analysis, model development, and hosting and drafting the manuscript. All the authors were involved in reviewing the manuscript and approved the final version of the manuscript.

### 7.6 Acknowledgments

We acknowledge the use of IBM Quantum Credits for this work. The views expressed are those of the authors and do not reflect the official policy opposition of IBM or the IBM Quantum team. We thank the administration and staff of R V College of Engineering for the support. We thank Suman A for her unwavering support.

## 8 Data and Software Availability statement

The tool can be accessed via GitHub at https://github.com/akshayuttarkarc/MotifQu and on PyPi via https://pypi.org/project/MotifQu/

