## Supplementary figures and images for "An inspectable Grover algorithm-based amplitude-amplification workflow for k-mer discovery and motif scanning with genomic coordinate traceability"

### genome_motifs.png

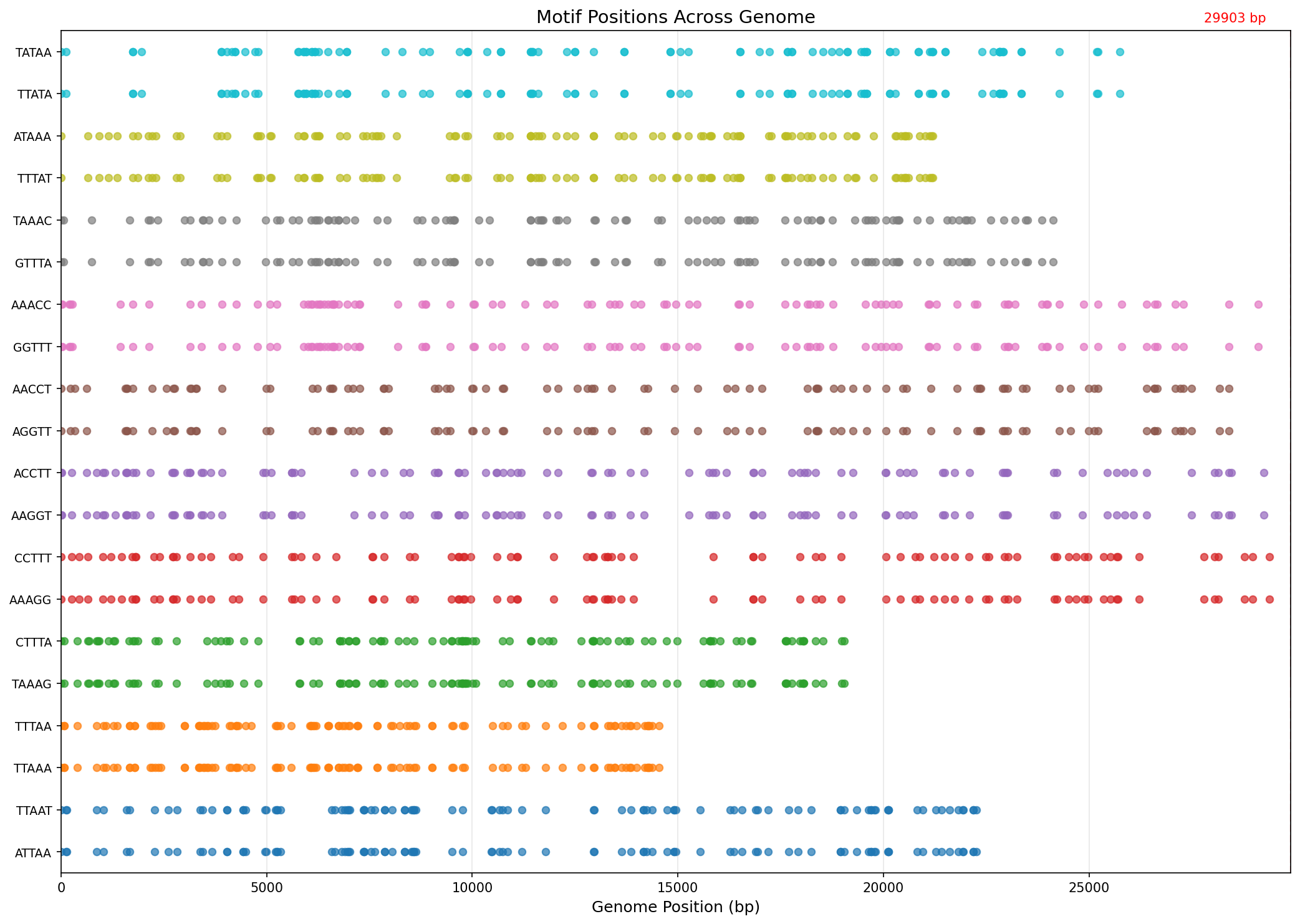

### genome_motifs.png

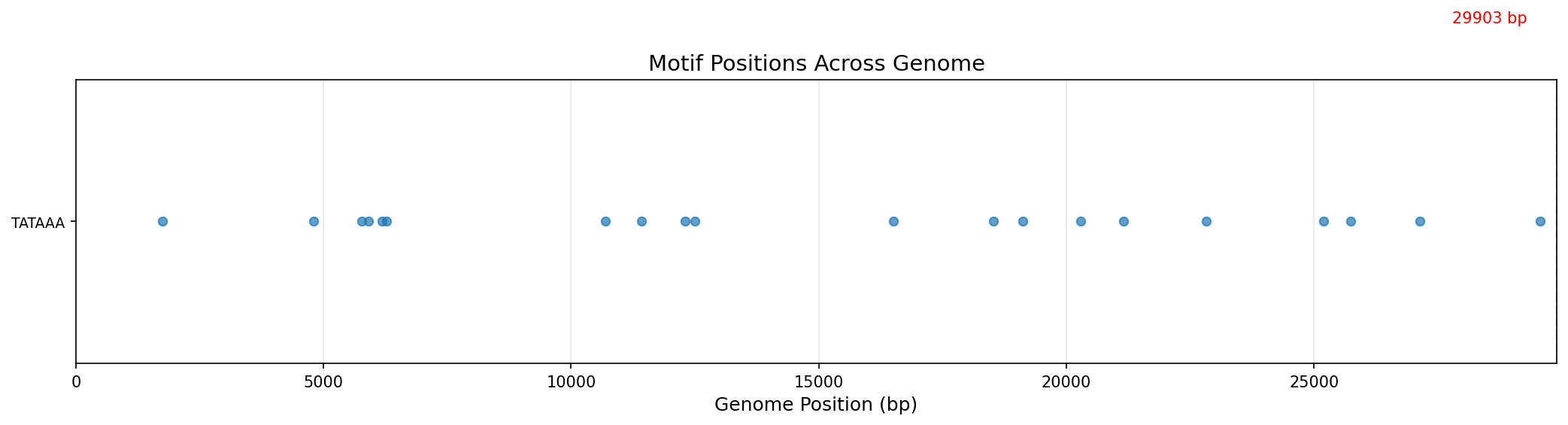

### genome_motifs.png

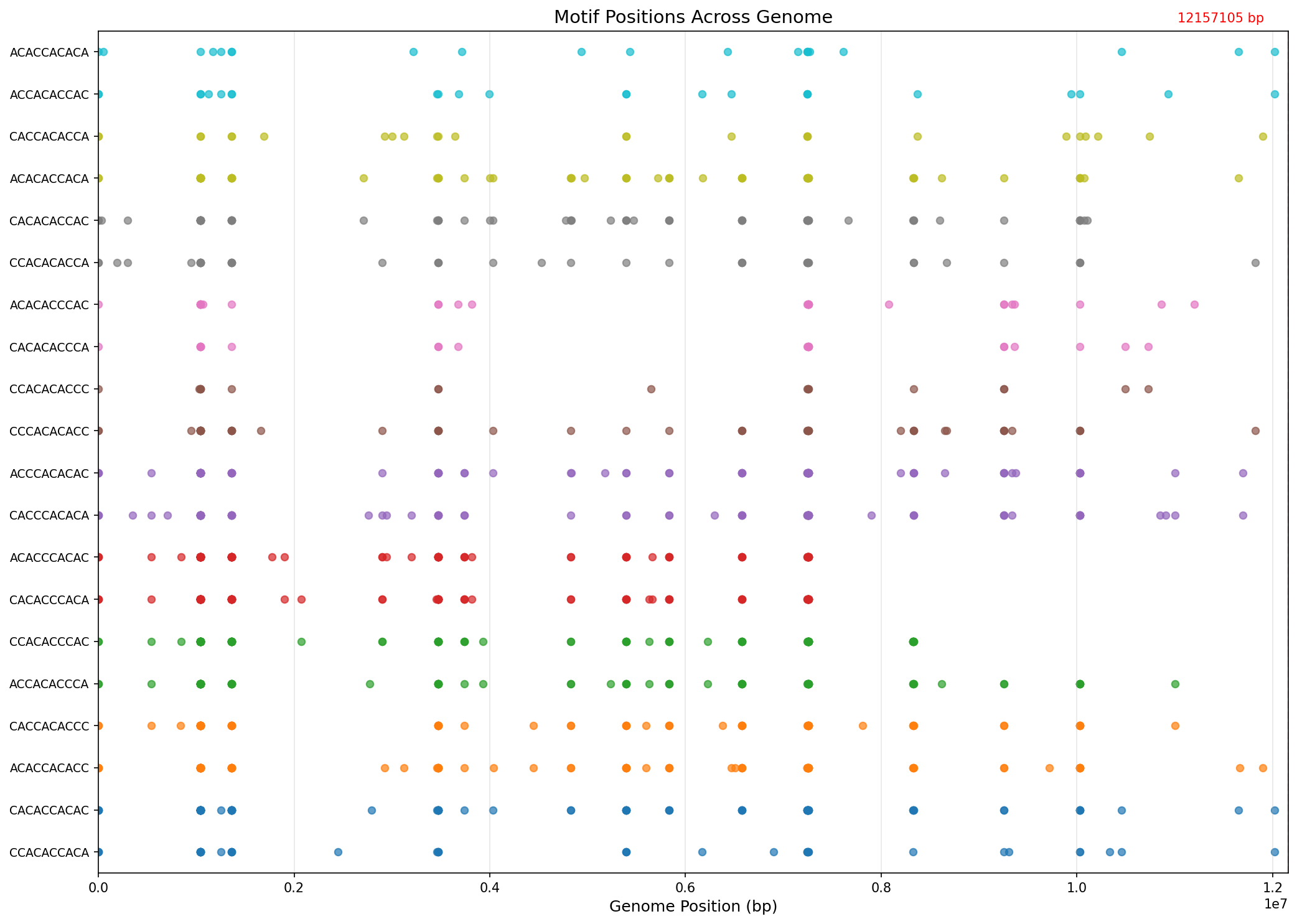

### grover_circuit.png

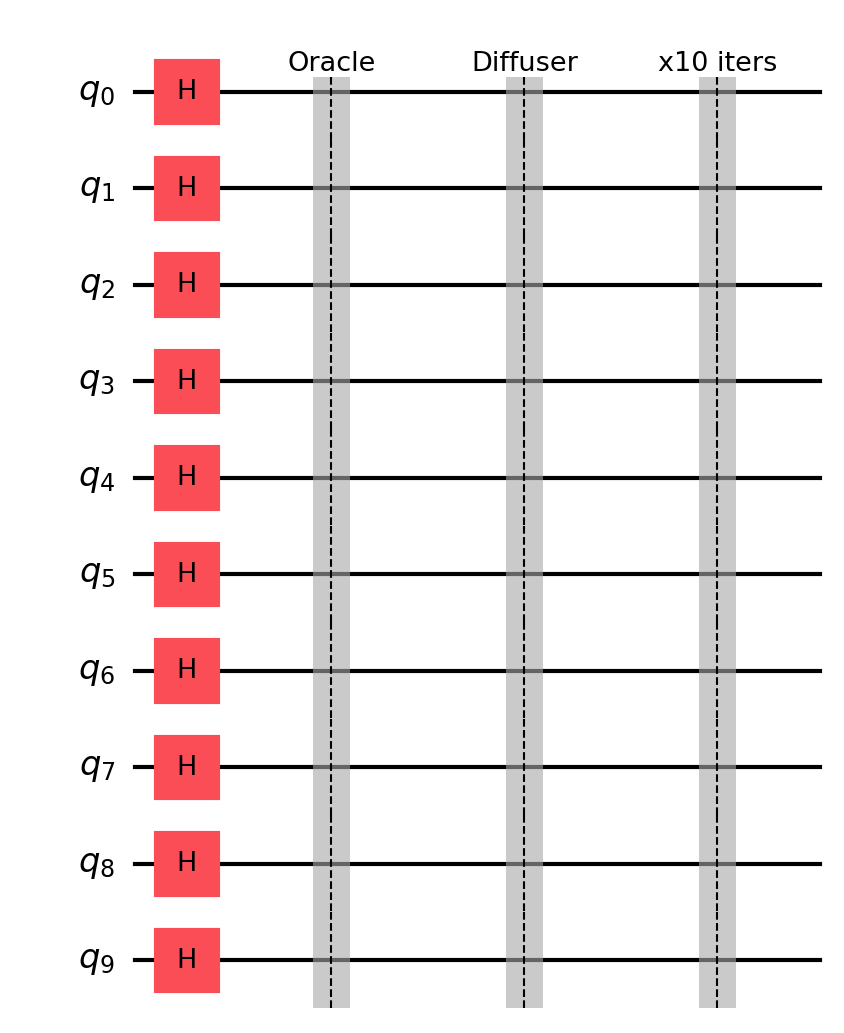

### grover_circuit.png

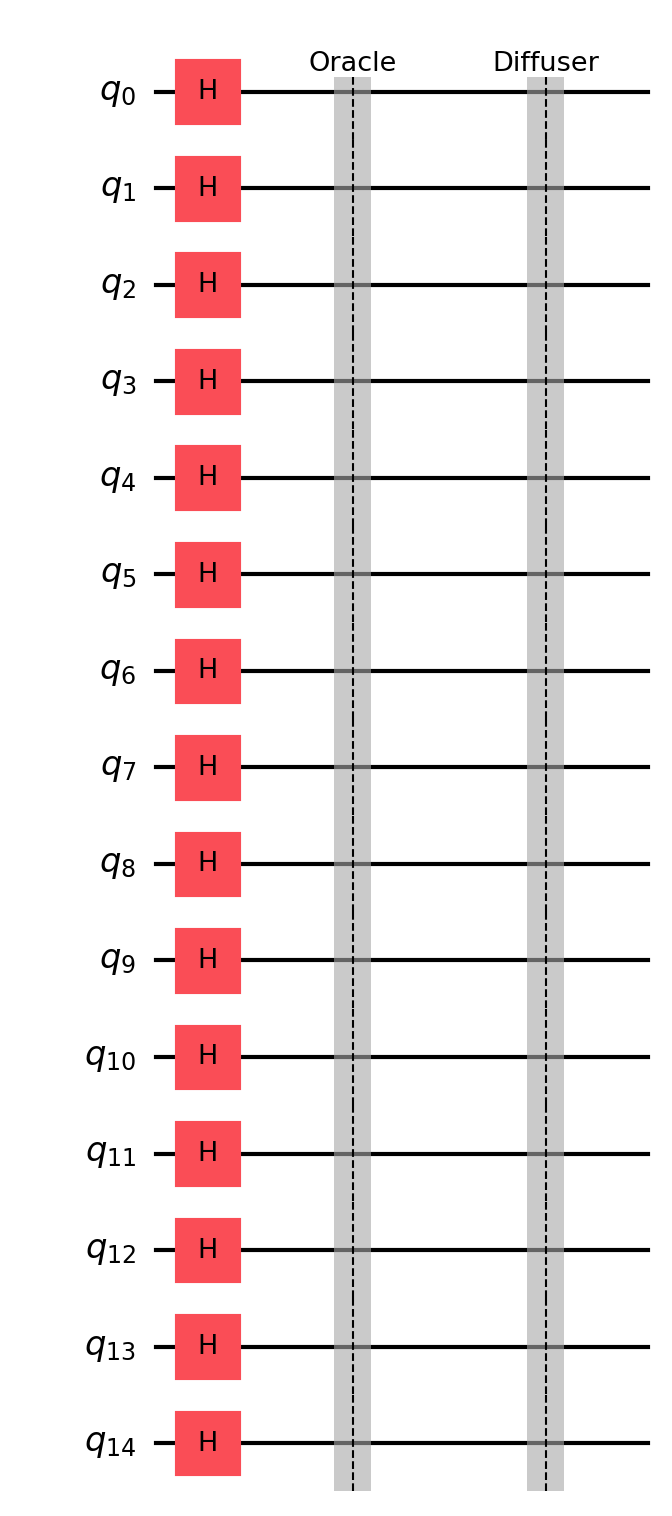

### grover_circuit.png

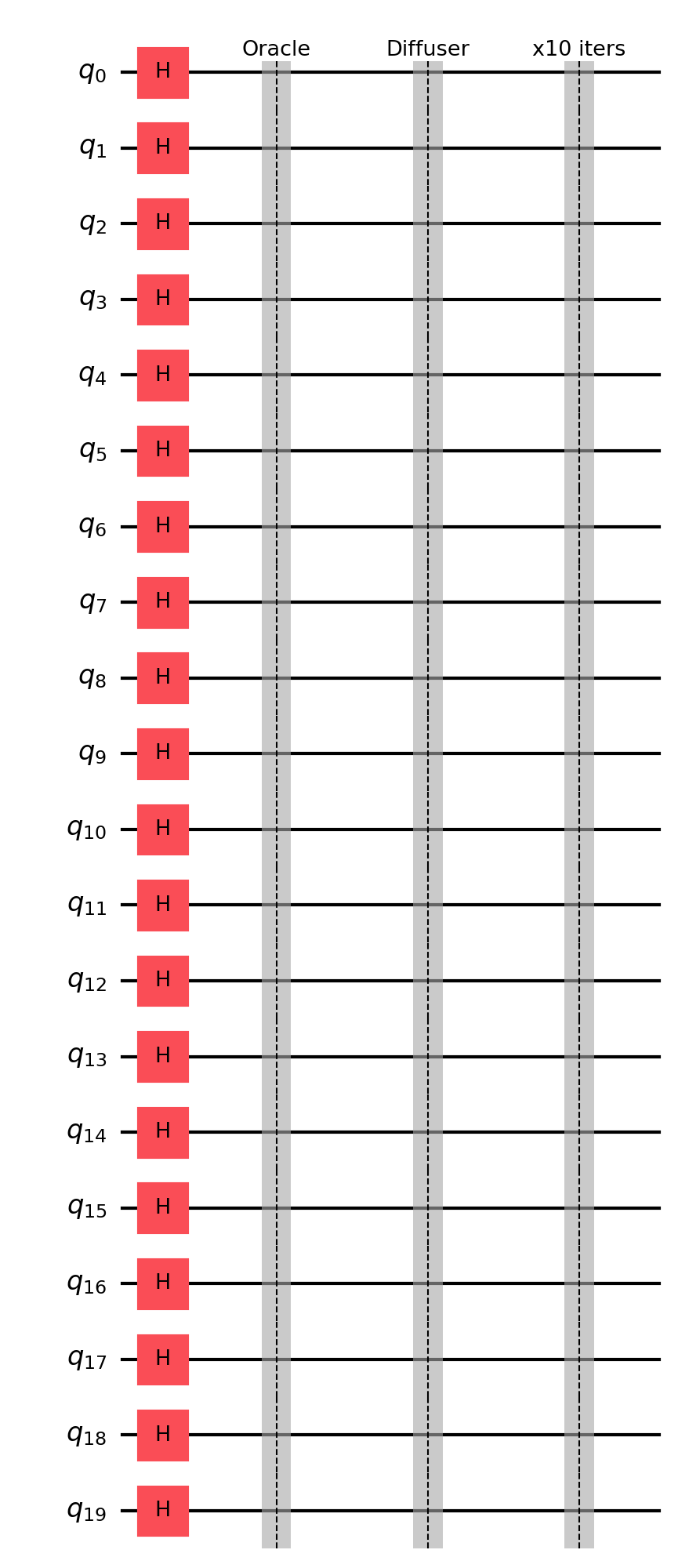

### job_d5terg8husoc73equlf0_results_meas.png

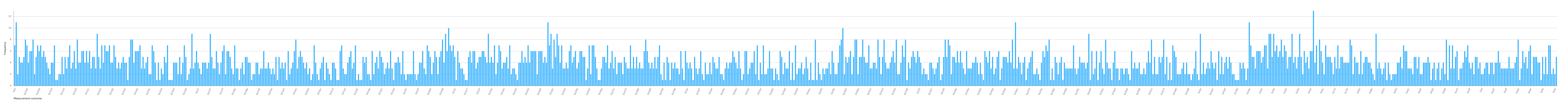

### job_d5ti5m8husoc73er3120_results_meas.png

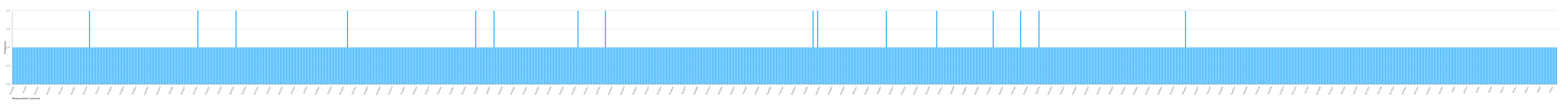

### probability_distribution.png

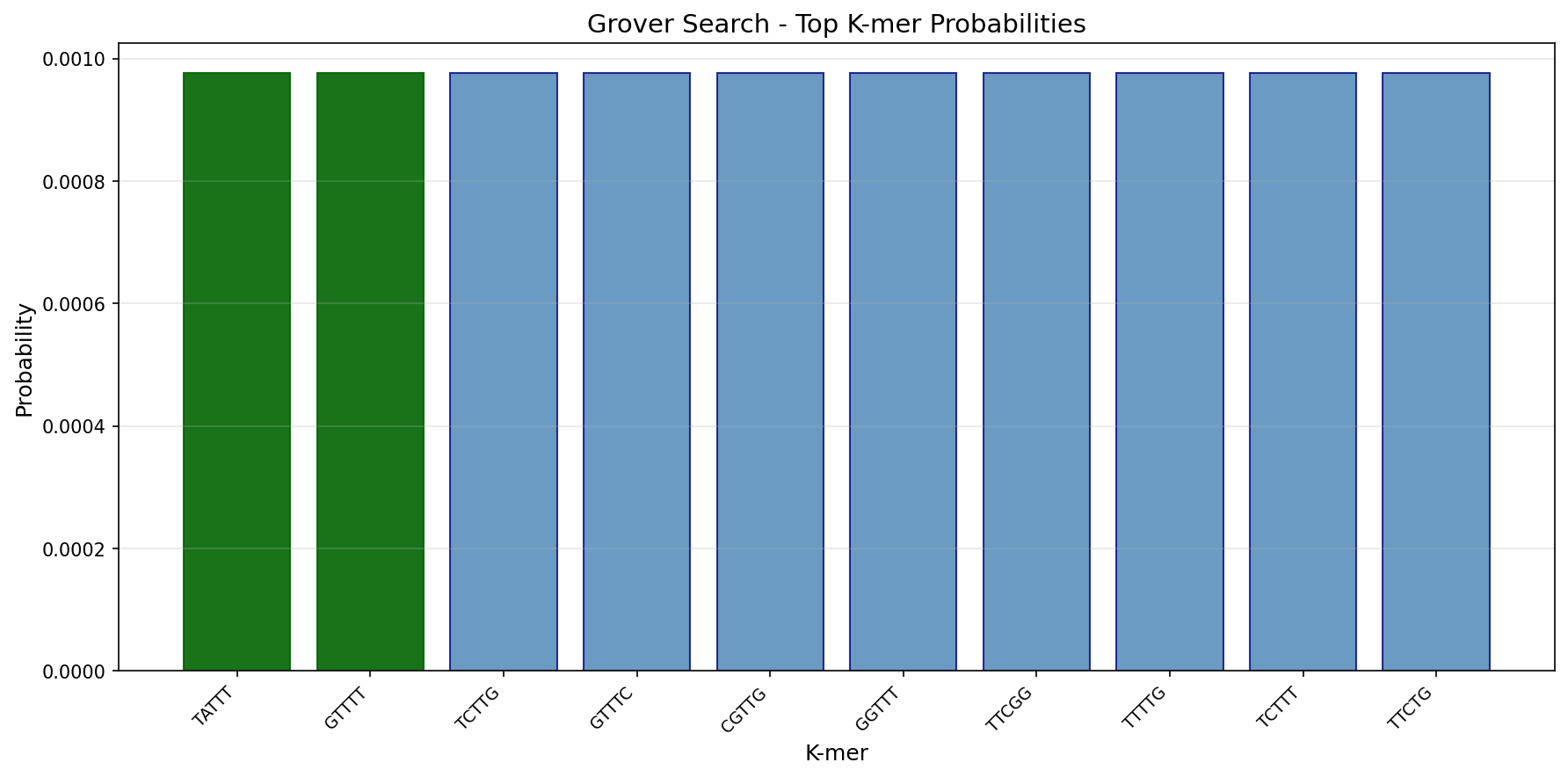

### probability_distribution.png

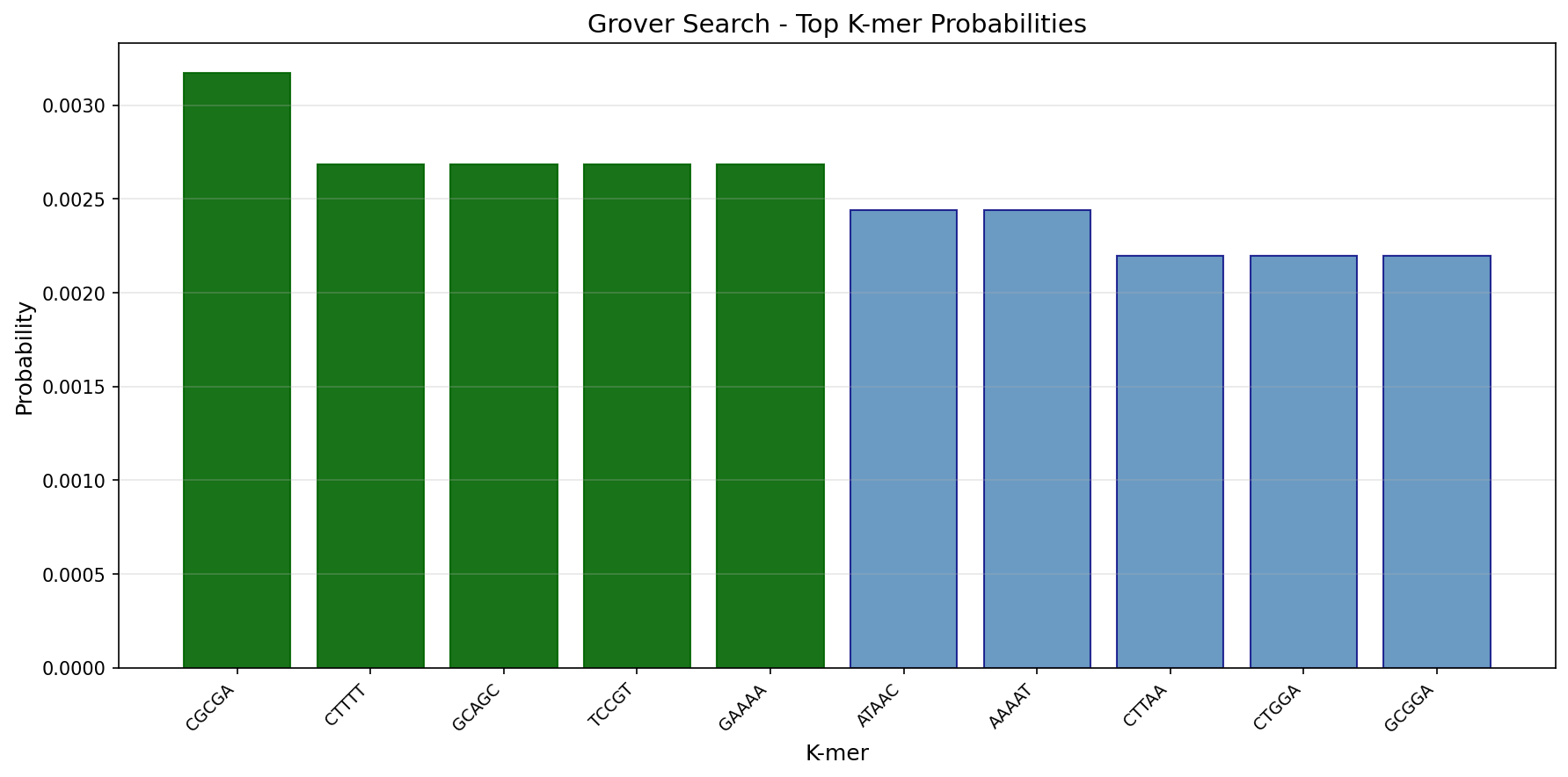

### probability_distribution.png

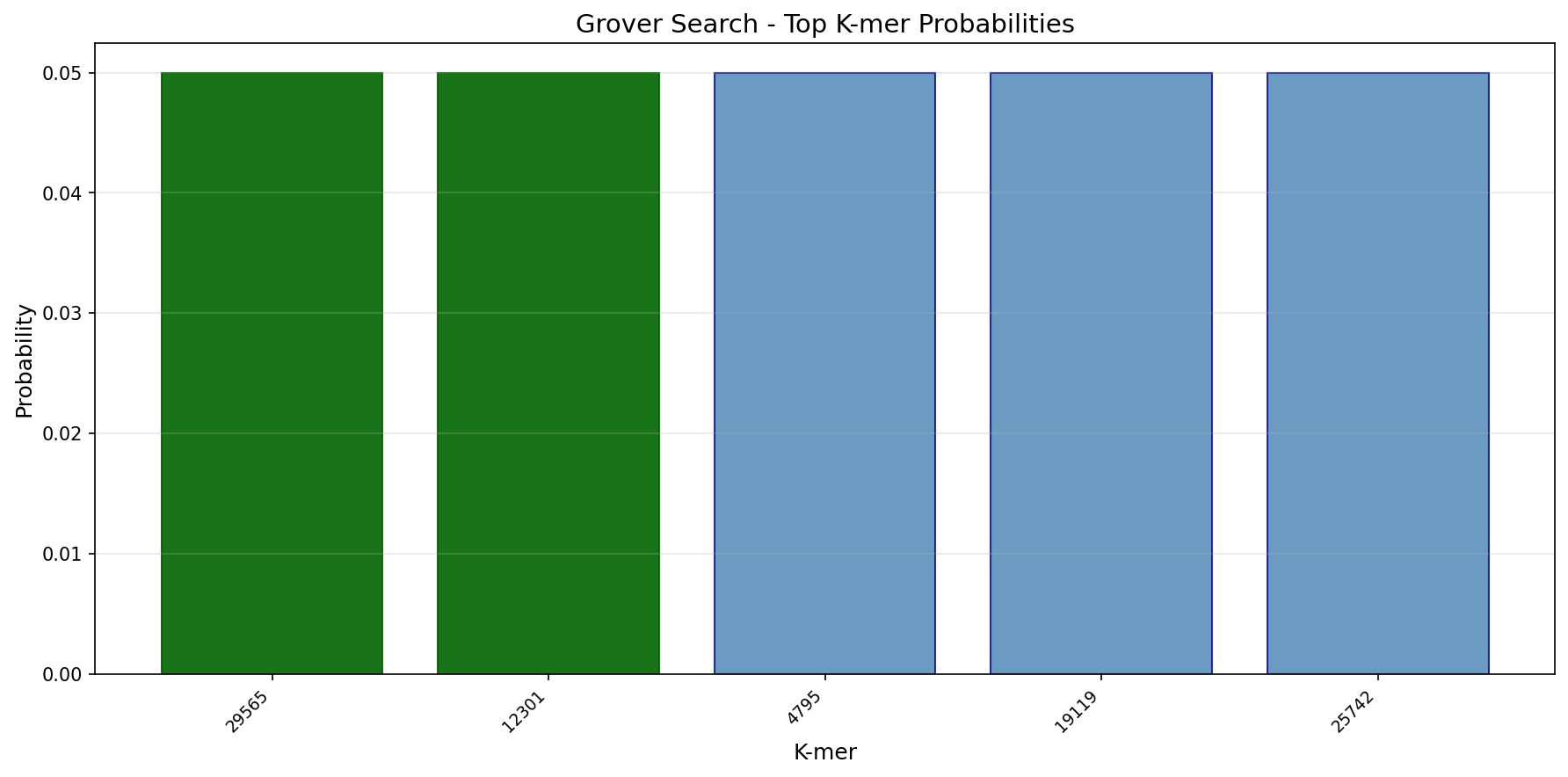

### probability_distribution.png

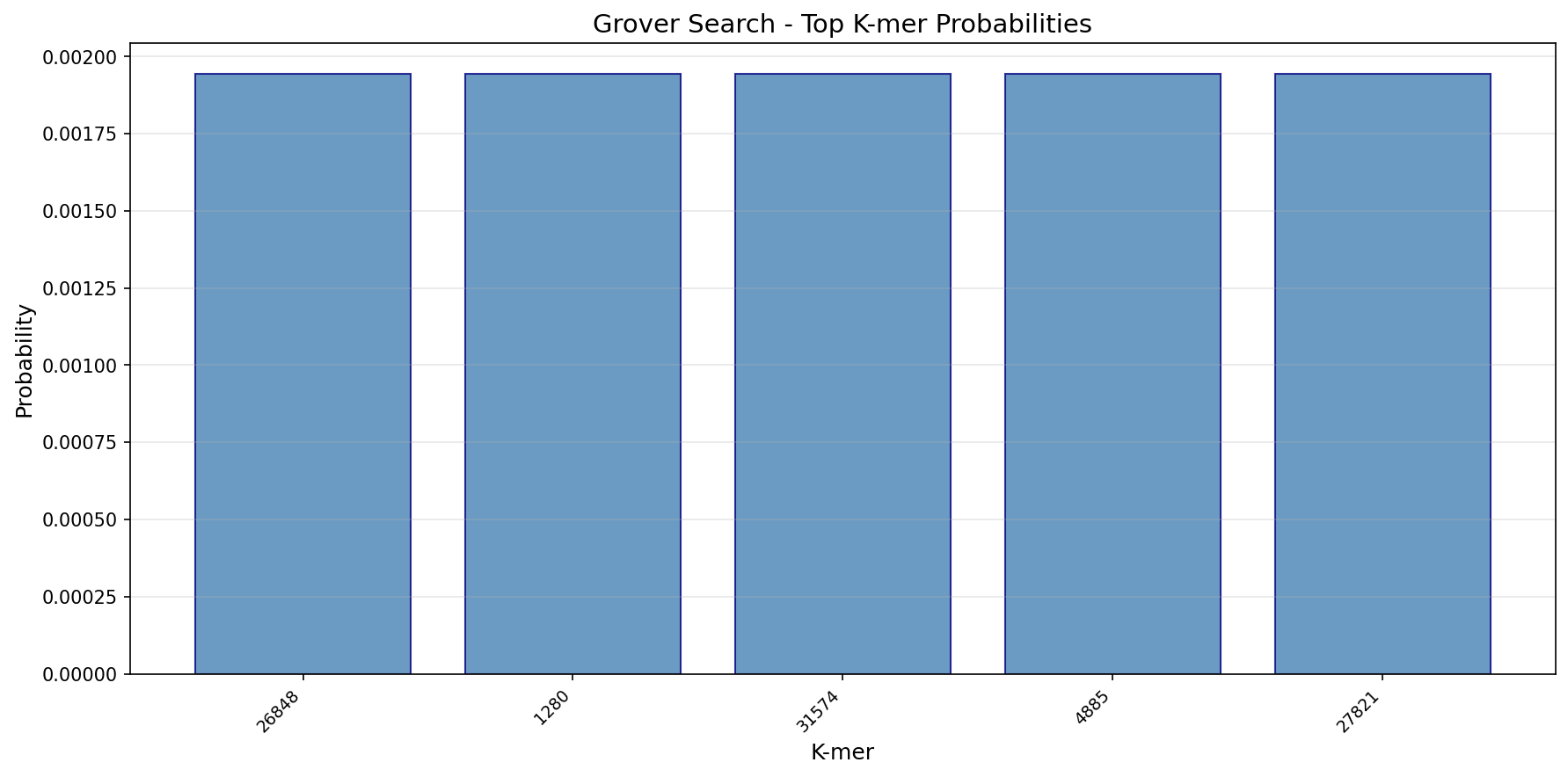

### probability_distribution.png

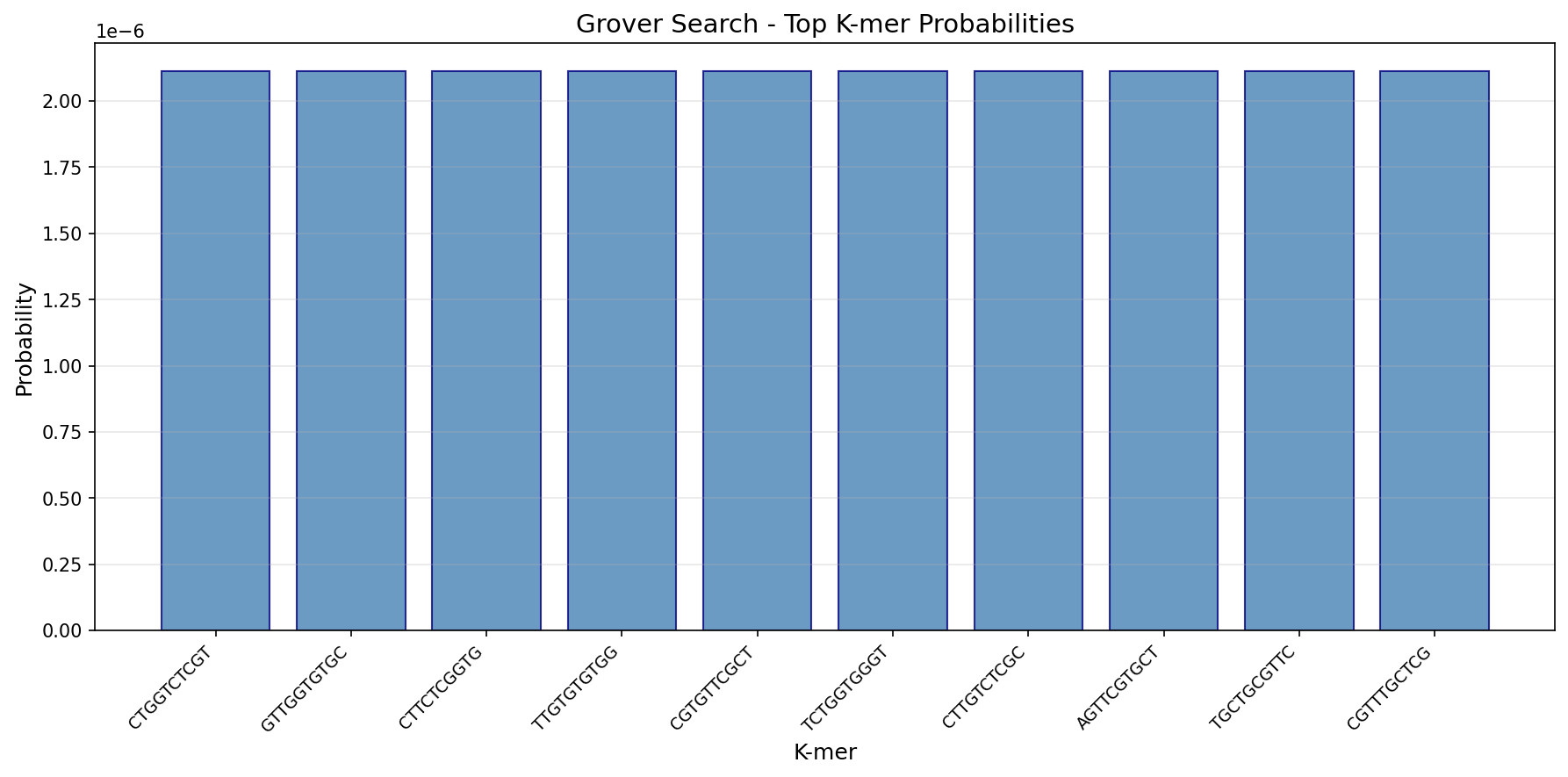
